# Erythrocyte TLR9 is upregulated in metabolic associated fatty liver disease, and is linked to an inflammatory immunometabolic signature

**DOI:** 10.1101/2024.01.23.576889

**Authors:** Charalampos Papadopoulos, Konstantinos Mimidis, Ioannis Tentes, Konstantinos Anagnostopoulos

**Affiliations:** Laboratory of Biochemistry, Department of Medicine, Democritus University of Thrace, Alexandroupolis, Greece; First Department of Internal Medicine and Laboratory for the Study of Gastrointestinal System and Liver, University Hospital of Alexandroupolis, Greece

**Keywords:** erythrocyte, MAFLD, TLR9, TSP-1, immunometabolism

## Abstract

Metabolic associated fatty liver disease (MAFLD) consists of lipid accumulation in the liver. Lipotoxicity, supported by aberrant amino acid metabolism, induces TLR9 upregulation, and activation, driving inflammation. Relatively, erythrocyte TLR9 activation leads to membrane rearrangement, surface CD47 loss, and pro-inflammatory erythrophagocytosis. Erythrocyte surface protein loss is accompanied by chemokine release. In addition, CD47 binds circulating TSP-1, a molecule controlling arginine and glutamine metabolism, along with metabolic inflammation. Based on these, we speculated that in MAFLD, lipotoxicity would drive erythrocyte TLR9 upregulation and activation, leading to immunometabolic remodeling.

Twenty-four patients (15 men and 9 women) with MAFLD and 9 healthy controls (4 men and 5 women) were enrolled. Erythrocytes were isolated from EDTA-containing blood. Protein levels were measured in erythrocyte lysates (triton X-100 0.01% v/v) or plasma with enzyme-linked immunosorbent assays, whereas lipids and enzyme activities were measured in erythrocyte hemoglobin-free membranes by a semi-quantitative thin layer chromatography and assay kits, respectively.

The levels of TLR9 were increased (p=0.002) and positively correlated with sphingosine levels, albeit not statistically significantly (p=0.060). The erythrocyte membrane PC/PE ratio was decreased (p=0.002) and inversely correlated to TLR9 levels. Erythrocyte TLR9 levels correlated inversely with CD47, and positively with MCP-1 release. TSP-1 was decreased in MAFLD erythrocytes (p=0.0017) and correlated positively with CD47 and negatively with TLR9 levels. In vitro, erythrocytes of MAFLD patients bound less TSP-1 molecules. Erythrocytes of MAFLD patients also exhibit decreased arginase-1 protein (p=0.009) and activity (p=0.042). Glutaminase activity was increased, albeit to not statistically significantly different levels (p=0.25). The levels of CD35 were not different (p=0.65), excluding the role of erythrocyte aging for explaining these events.

In MAFLD patients, erythrocyte TLR9 is upregulated and is associated with erythrocyte sphingolipid and glycerophospholipid perturbation, CD47 loss, MCP1 release, reduced TSP-1 scavenging, decreased arginase and increased glutaminase.

## INTRODUCTION

The pandemic of obesity constantly progresses worldwide and impacts the quality of life and economy of millions of people^1^. During obesity, the excess fatty acids triggers a pro-inflammatory phenotype of adipose tissue and leads to triglyceride storage in non-adipose tissues, such as skeletal muscle, pancreas, and liver. Eventually, due to the limited storage capacity of these tissues, toxic lipids are formed such as ceramide, sphingosine 1-phosphate, lysophosphatidylcholine etc. These lipid classes lead to lipotoxicity^2^. Lipotoxicity drives organelle dysfunction, cell death, inflammation and fibrosis^3^. The umbrella of these mechanisms when taking place in the liver constitutes non-alcoholic fatty liver disease (MAFLD). Mechanistically, hepatic lipotoxicity, is preceded by aberrant arginine and glutamine metabolism^4,5^. Next, lipotoxicity leads to TLR9 upregulation in the liver^6–8^ and mitochondrial DNA release which activates TLR9^8–11^. In fact, these events mediate the transition from hepatic steatosis to inflammation^7,9^ and fibrosis^12^. Similarly, limited T-cell TLR9 expression protects from hepatic metabolic inflammation^13^. Although the mechanisms linking lipotoxicity to TLR9 upregulation are diverse, there is evidence that sphingosine-induced cholesterol accumulation^14^ dampens TLR9 degradation^15^.

Despite the extended research on MAFLD, erythrocytes are rarely explored. Apart from gas exchangers, erythrocytes also possess central role in the immunometabolic regulation^16,17^. Accordingly, erythrocytes exhibit extended lipidome rearrangement, removal signals and pro-inflammatory phagocytosis during MAFLD^18–22^.

Importantly, recent studies have unveiled that erythrocyte TLR9 ligation is necessary for mitochondrial DNA-induced inflammation through pro-inflammatory erythrophagocytosis^23^. On a molecular level, erythrocyte TLR9 activation triggers membrane rearrangement through lipid remodeling, and CD47 loss^23^. Of note, surface protein loss is also followed by chemokine release^24^. In addition, erythrocyte surface CD47 binds TSP-1 and regulates erythrocyte arginine metabolism^25^. TSP-1 also regulates the metabolism of glutamine^26^, which partially explains why TPS-1 participates in metabolic inflammation in the liver^27^.

Based on these, we speculated that lipotoxicity in MAFLD would result in erythrocyte TLR9 upregulation and activation. As such, erythrocyte TLR9 would be increased and would correlate with erythrocyte sphingosine, glycerophospholipids levels, CD47 loss, MCP-1 release, TSP-1 binding, arginase and glutaminase levels.

## 2. Materials and Methods

### SUBJECTS

Twenty-four patients (15 men and 9 women) with MAFLD and 9 healthy controls (4 men and 5 women) were enrolled. All healthy controls were nonobese (BMI<25), free of chronic diseases, had no signs of infection in the previous 2 months, and had normal AST and ALT levels. They were selected from the outpatient liver clinic of the First Department of Internal Medicine, School of Medicine, Democritus University of Thrace, Alexandroupolis, Greece. Hepatic steatosis was assessed by conventional ultrasound performed by a skilled radiologist according to the European guidelines for adults at risk for MAFLD. All patients had elevated AST and ALT levels at baseline. Viral hepatitis (A, B, C, D, E) was ruled out by ELISA and/or polymerase chain reaction test. Alcoholic and drug induced hepatitis was excluded by a questionnaire, and autoimmune hepatitis by antibody detection. Liver fibrosis stage was also evaluated by noninvasive serum markers (Fibroscan, FIB-4, AST/ALT ratio). Table 1 shows their anthropometric and clinical characteristics. The Scientific Council of the University Hospital of Alexandroupolis and the Ethics Committee approved the study after obtaining informed consent from the participants.

**Table 1.**
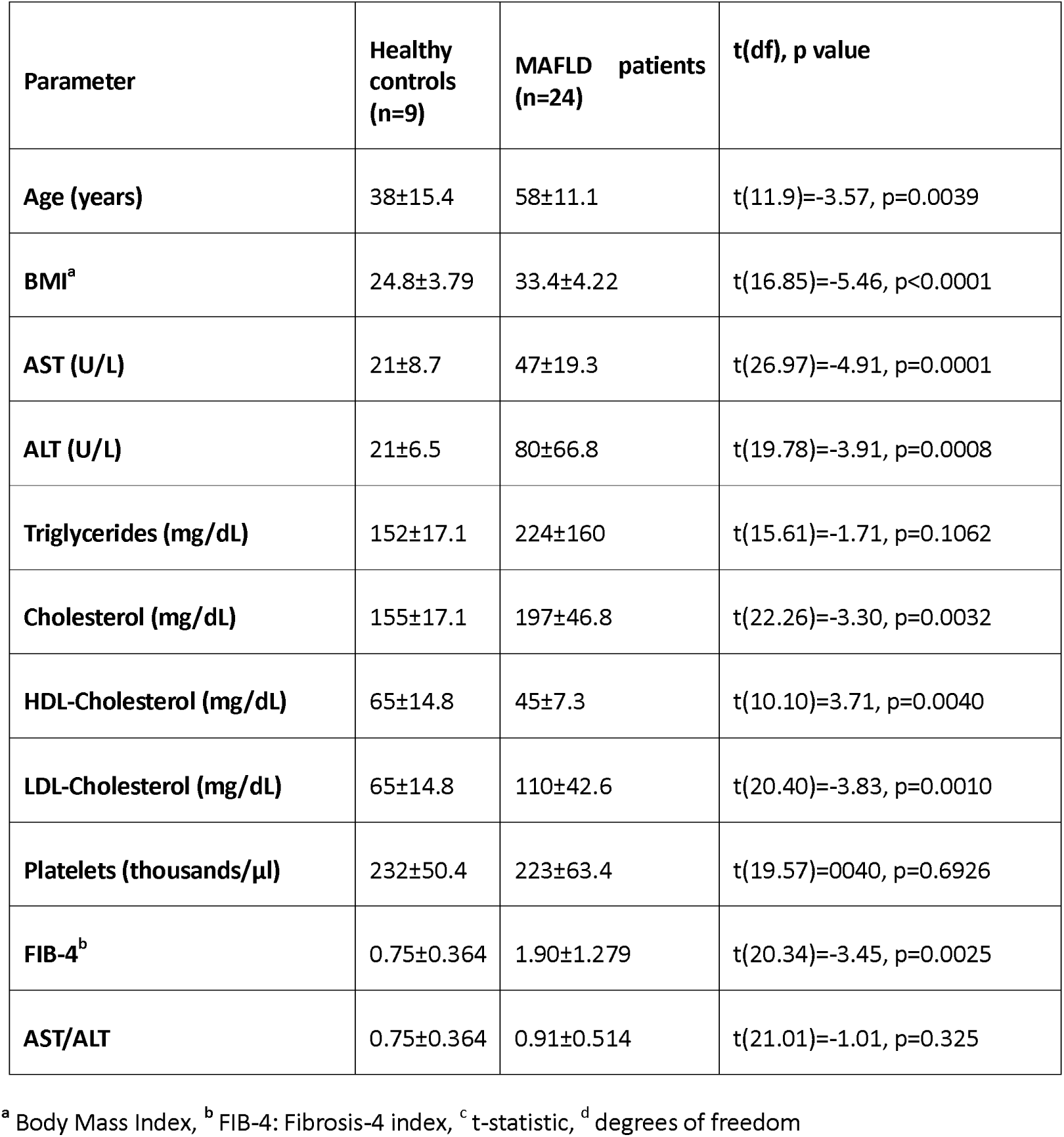
Anthropometric and clinical characteristics of NAFLD patients and healthy subjects. Results are reported as mean±Standard Deviation. The p-value is for the difference between means.

#### Erythrocyte isolation

From every patient or healthy volunteer three ml of blood in tubes containing 5.4 mg ethylenediaminetetraacetic acid (EDTA) were isolated. To isolate erythrocytes, the tubes containing blood were centrifuged at 200×g for 10 min at 4C in order to remove the buffy coat and plasma. Then, 1 mL of erythrocyte pellet was washed with cold saline solution and centrifuged at 200×g for 10 min at 4C. The last step was repeated four times, until a clear erythrocyte population was achieved.

#### Erythrocyte membrane isolation

After erythrocyte isolation, 500 μl of packed erythrocytes were incubated for 30 min at 4°C with continuous shaking, diluted with cold hemolysis solution (Tris 1 mM-NaCl 10 mM-EDTA 1 mM, pH 7.2 at a final concentration of 1:10 (vol/vol)). This was followed by centrifugation at 10,000×g, at 4°C for 15 min. The pellet, corresponding to erythrocyte membranes, was collected. The last step was repeated as many times as needed, until the removal of hemoglobin-free erythrocyte membranes. Samples were kept at −80°C until analysis.

#### Lipid extraction

For lipid extraction, we use a modified Folch method. ^28^ Briefly, 500 μL of 2:1 chloroform/methanol was added to 100 μL of erythrocyte membrane. was After a vortexing for 30 sec and a centrifugation for 10 min at 5000×g, the organic phase was collected. The previous step was repeated for the aqueous phase. The two organic phases were washed with water to remove any remainders of polar molecules by centrifugation for 10 min at 5000×g and then evaporated at 70°C.

#### Lipid separation and visualization

Thin-layer chromatographic (TLC) analysis of lipids was performed on chromatographic plates (TLC Silica gel 60 F254; Merck KGaA 64071, Darmstadt, Germany) according to our previously published protocol^20^.

#### Phosphatidylcholine and Phosphatidylethanolamine quantitation

The method for PC and PE quantitation is based on the intensity of the green color of the band corresponding to PC or PE, respectively. The characteristics of the method (specificity, linearity, accuracy, precision, limit of detection, limit of quantification and sensitivity) can be found in our previous study^20^.

#### Production of erythrocyte conditioned media

Erythrocytes were incubated with RPMI at concentration of 10^7^ erythrocytes/ml, for 4 hours, with recombinant TSP-1 (Origene, USA) at a concentration of 100ng/ml. TSP-1 was diluted in PBS. After incubation, conditioned media were isolated with centrifugation at 200g for 10 minutes.

#### Lysis of erythrocytes

One milliliter of packed erythrocytes was lysed with Triton X-100 at a final concentration of 0.01% vol/vol ^21^.

#### Determination of arginase, TLR9, CD35 and TSP-1 protein levels

The levels of TLR9, CD35, TSP-1 and arginase-1 were determined by ELISA in erythrocyte lysates, conditioned media or plasma according to the manufacturer’s instructions (OriGene, USA).

#### Determination of erythrocyte arginase-1 and glutaminase-1 activity

The activity of erythrocyte arginase-1 and glutaminase-1 was determined in erythrocyte membrane preparations according to the manufacturer’s instructions (Fine-Test, China and Abbexa. UK, respectively).

#### Statistical analysis

Results were expressed as mean ± standard deviation and median, unless otherwise stated. Statistical analysis was performed with the R programming language v. 4.0^29^ Welch’s independent samples t-test^30^ was used for comparison of means between groups. This test is an adaptation of Student’s t-test, and does not assume equal variances and/or sample sizes, being more reliable when these conditions occur. Glass’s Δ was used to quantitate the effect size^31^ since it is more appropriate when the standard deviations between groups are significantly different. According to the above, effect sizes with values Δ<0.2 are considered very small, small for 0.2≤Δ<0.41, moderate for 0.41≤Δ<0.63 and large for Δ≥0.63.

The results are reported as t(degress of freedom)= test statistic, the p-value, followed by Glass’s Δ (along with is 95% confidence interval, CI), the statistical power 1-β for a significance level α=0.05 and the 95% confidence interval for the difference between means of the healthy and MAFLD groups, as shown in Table 2.

**Table 2.**
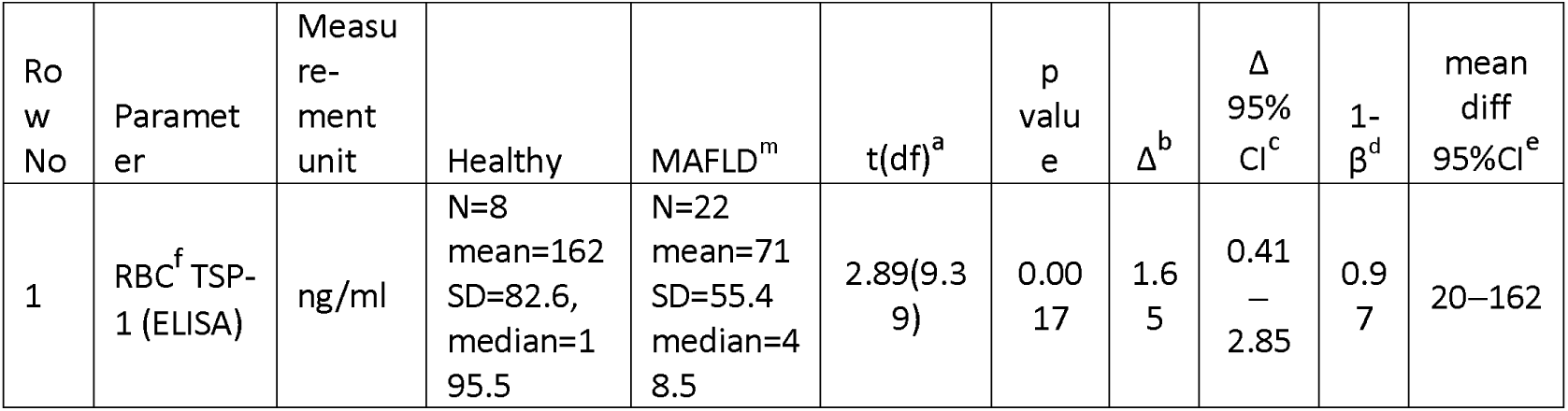
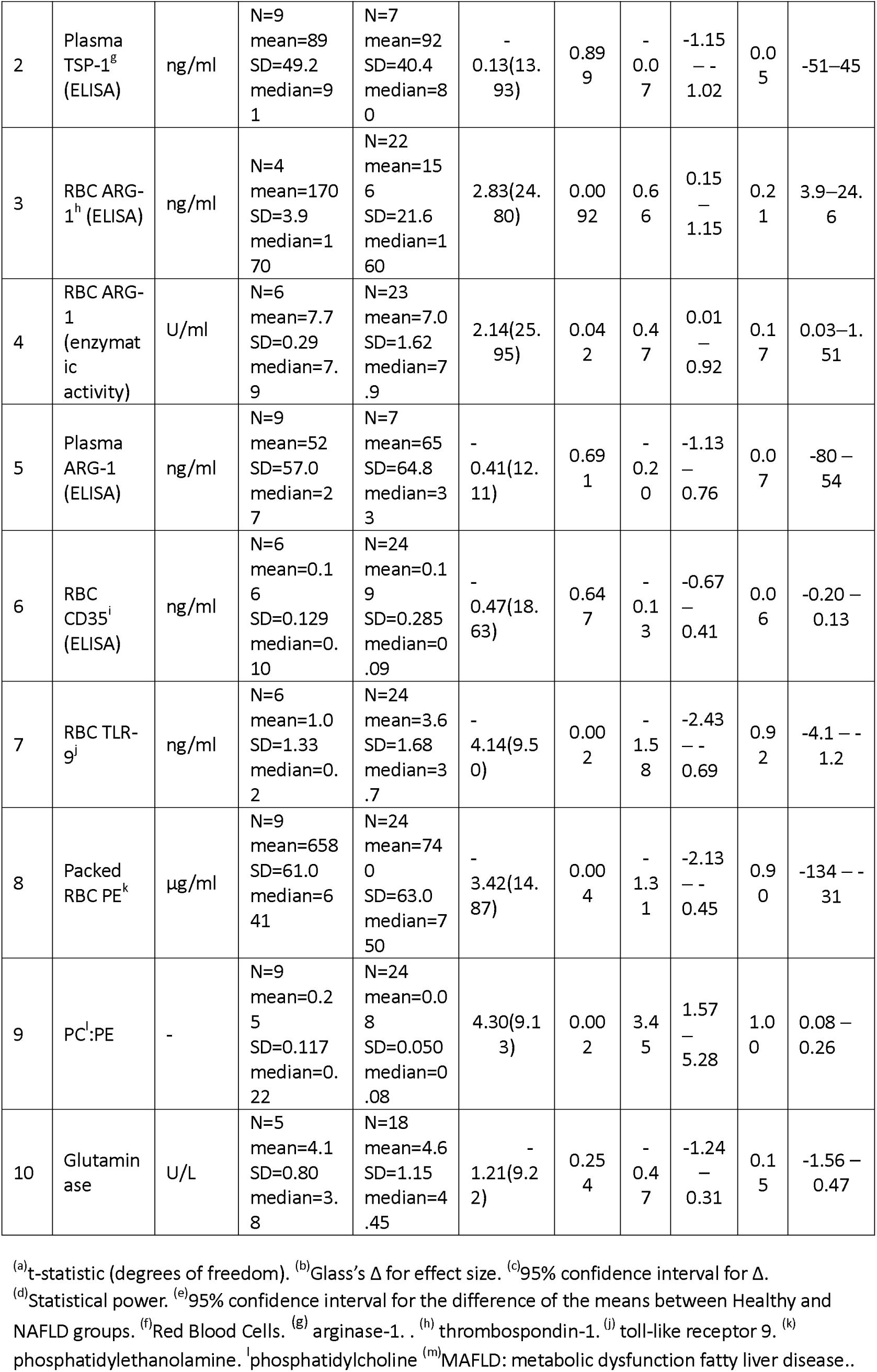
Test for difference between means using Welch’s t-test.

Linear regression was performed using the base R lm command. All plots were created using the ggplot package. In all the boxplots, the box extends from the lower quartile to the upper quartile (that is the range corresponds to the inter-quartile range, IQR). The vertical lines extend to ±1.5×IQR. Individual points shown correspond to extreme values, outside the ±1.5×IQR.

## RESULTS

### ERYTHROCYTE TLR9 LEVELS

Erythrocytes express TLR9^23^. This receptor binds and carries mitochondrial DNA to macrophages after erythrophagocytosis, leading to inflammation^23^. In addition, TLR9 ligation can lead to CD47 loss^23^. For these reasons, we quantified TLR9 on erythrocytes. In MAFLD patients TLR9 levels were statistically significantly greater (mean=3.6 ng/ml) than in healthy subjects (mean=1.0 ng/ml), with p=0.002 and a large effect size (Table2 and Figure1).

**Figure 1:**
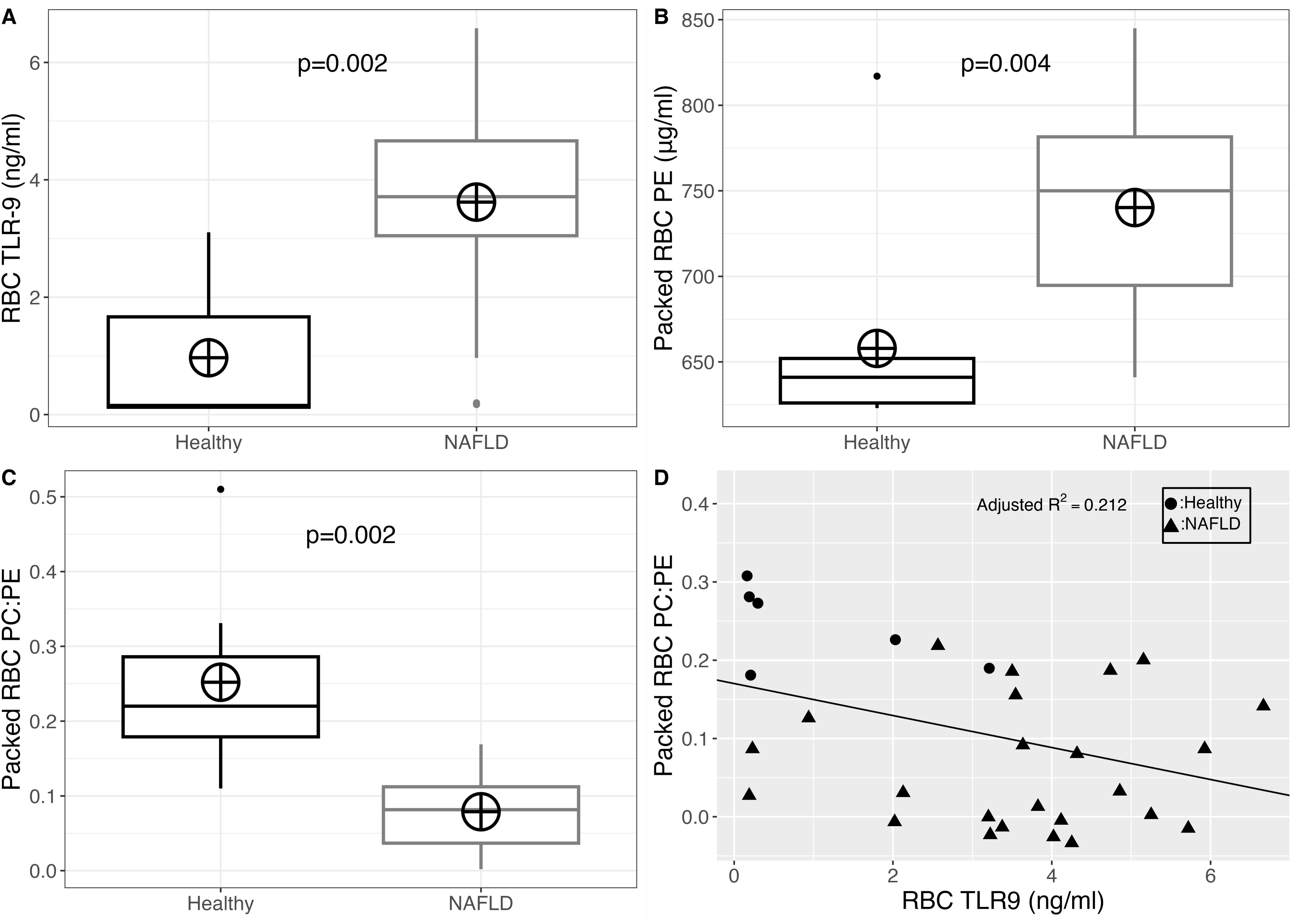
A. Erythrocyte TLR9 levels in healthy controls (n=6) and MAFLD patients (n=24). B. Erythrocyte phosphatidylethanolamine levels in healthy controls (n=9) and MAFLD patients (n=24). C. Erythrocyte phosphatidylcholine to phosphatidylethanolamine ratio in healthy controls (n=9) and MAFLD patients (n=24) D. Linear regression between erythrocyte TLR9 and erythrocyte PC/PE ratio in healthy controls (n=6) and MAFLD patients (n=24).; MAFLD: metabolic associated fatty liver disease; PC: phosphatidylcholine; PE: phosphatidylethanolamine; RBC: red blood cells; TLR9: toll-like receptor 9.

### ERYTHROCYTE TLR9 AND SPHINGOSINE CORRELATION

Lipotoxicity is known to drive TLR9 upregulation^6^. Because erythrocytes lack nucleus and endoplasmic reticulum, we hypothesized that the increased TLR9 levels could be explained by reduced degradation. Accordingly, previous studies have shown that sphingosine regulates protein trafficking and degradation^14,15^. Importantly, we have shown in the past sphingosine accumulation in erythrocytes of NALFD patients^21^. We found here that the levels of TLR9 correlate with sphingosine (r=0.94, p=0.603)(supplementary figure 1). However, the limited number of subjects for these measurements hinders this association.

### ERYTHROCYTE MEMBRANE PHOSPHATIDYLETHANOLAMINE LEVELS AND PC/PE RATIO

Erythrocyte TLR9 activation increases the surface TLR9 content, possibly through lipid remodeling^23^. We hypothesized that the phosphatidylcholine to phosphatidylethanolamine ratio would be correlated to TLR9. There was a statistically significant elevation in the PE levels of packed erythrocytes between MAFLD patients (mean=740 μg/ml) and healthy subjects (mean=658 μg/ml), with p=0.004 and a large effect size (Table 2 and Figure Figure1). In addition, the PC/PE ratio decreased (healthy mean=0.25, MAFLD mean=0.08), with p=0.02 and a large effect size, as shown in Table 2 and Figure1

We showed that TLR-9 levels are inversely correlated with the PC/PE ratio. When linear regression was performed, the adjusted R^2^ was 0.212, F(1,28)=8.779 (Figure1).

### CORRELATION BETWEEN TLR9 AND CD47

CD47 levels (quantified in our previous study^21^) were inversely correlated with TLR9, with an adjusted R^2^=0.177, F(1,21)=5.717. The intercept was 2548.4 (SE=545.1, p=0.0001) and the slope was −338.7 (SE=141.6, p=0.026) (Figure 2)

**Figure 2:**
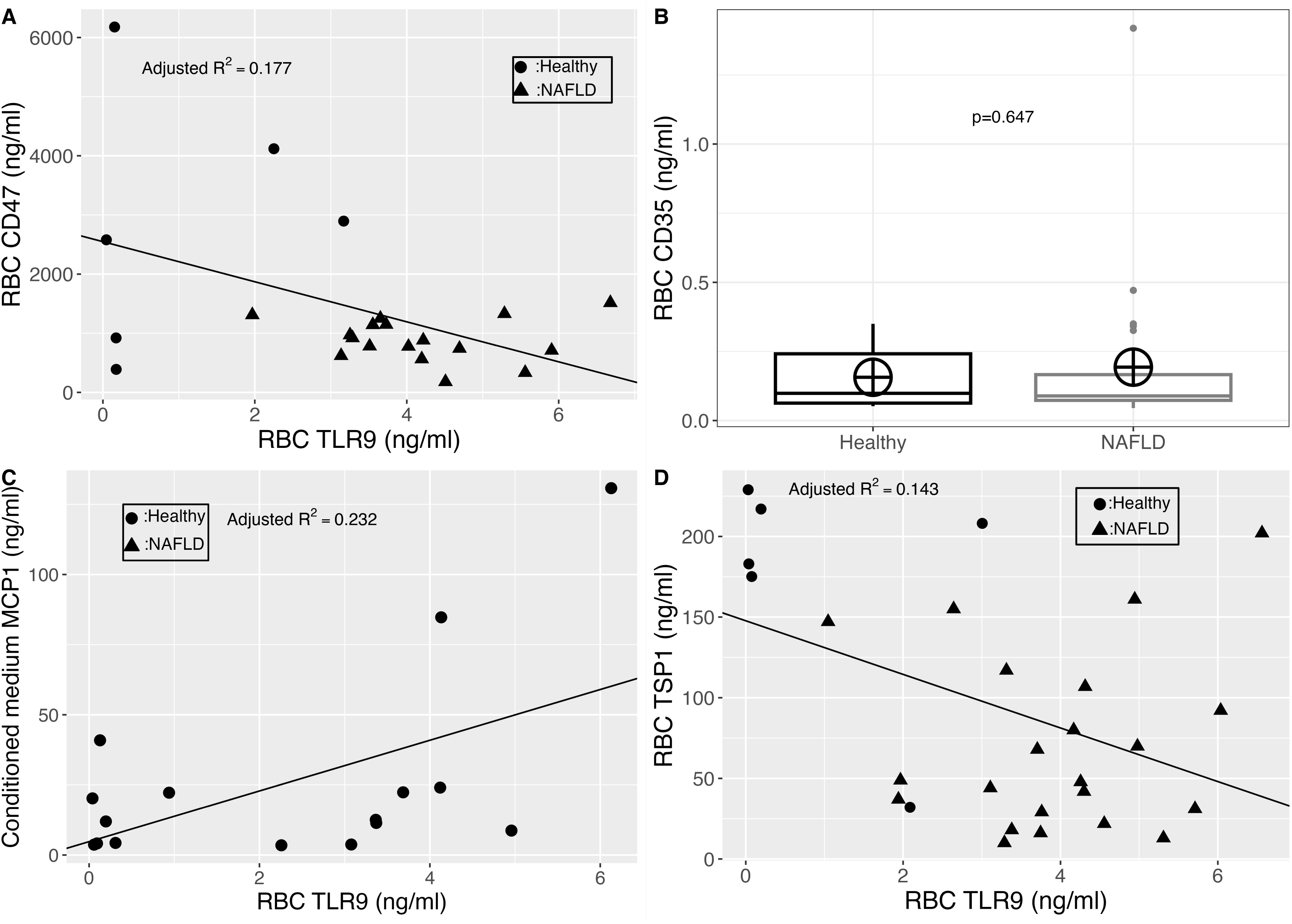
A. Linear regression between erythrocyte TLR9 and erythrocyte CD47 levels. B. Erythrocyte CD35 levels in healthy controls (n=6) and MAFLD patients (n=24). C. Linear regression between erythrocyte TLR9 and erythrocyte MCP-1 levels. D. Linear regression between erythrocyte TLR9 and erythrocyte TSP-1 levels. CD47: cluster of differentiation 47; CD35: cluster of differentiation 35;; MAFLD: metabolic associated fatty liver disease; RBC: red blood cells; TPS-1: thrombospondin-1;TLR9: toll-like receptor 9.

### CORRELATION BETWEEN TLR9 AND MCP-1

Erythrocyte surface protein loss is accompanied by chemokine release^24^. Based on the correlation between erythrocyte TLR9 and CD47, we speculated that TLR9 levels would be inversely correlated with MCP-1 release (measured in our previous study^22^). We found that erythrocyte TLR9 levels are inversely correlated with the release of MCP-1 in vitro (r=0.48, p=0.03)(figure 2).

### ERYTHROCYTE CD35 LEVELS

Erythrocyte CD47 loss is also related to erythrocyte aging^32^, and CD35 levels have been shown in the past to be a marker of erythrocyte aging^33^. For these reasons, we also quantified erythrocyte CD35. There was no statistically significant difference between patients and healthy controls. In healthy subjects the mean was 0.16 ng/ml and in MAFLD patients it was 0.19 ng/ml, with p=0.647 and a very small effect size (figure 2).

### ERYTHROCYTE TSP-1 LEVELS

Based on our previous results of reduced erythrocyte CD47 levels in MAFLD patients^21^, we examined the levels of bound TSP-1 on erythrocytes. On average, MAFLD patients had lower levels of erythrocyte TSP-1 (mean=71 ng/ml), than healthy subjects (mean=162 ng/ml). This difference was statistically significant (p=0.0017), and with a large effect size, as shown in Table 2, and Figure3.

### PLASMA TSP-1 LEVELS

Since we observed reduced levels of bound TSP-1 on erythrocytes of MAFLD patients, we wondered whether this difference is dependent on changes on the plasma levels. As shown in Table2 and Figure3, there was no statistically significant difference in plasma TSP-1 between MAFLD patients (mean=92 ng/ml) and healthy subjects (mean=89 ng/ml). Hence, the reduced levels of TSP-1 on erythrocytes of MAFLD patients are not explained by reduced levels in the plasma.

### CORRELATION BETWEEN ERYTHROCYTE CD47 AND TSP-1 LEVELS

Since the levels of erythrocyte-bound TSP-1 seem to be largely independent of the plasma levels, we hypothesized that alterations on erythrocytes are responsible for the observed difference. Previous studies have shown that TSP-1 binds CD47 of erythrocytes^34^. Linear regression was performed with erythrocyte CD47 as the independent and TSP-1 as the dependent variable. The scatterplot along with the resulting regression line is shown in Figure 3. The adjusted R^2^ was 0.227, F(1,23)=8.04, p=0.0093. The intercept was 58.5 (SE=19.03, p=0.0054) and the slope was 0.023 (SE=0.0082, p=0.0094). Since the adjusted R^2^ is 0.227, this means that 22.7% of the variation in RBC TSP-1 can be explained by the variation in RBC CD47. Erythrocyte TSP-1 levels were also inversely correlated with TLR9 levels (figure 2).

**Figure 3:**
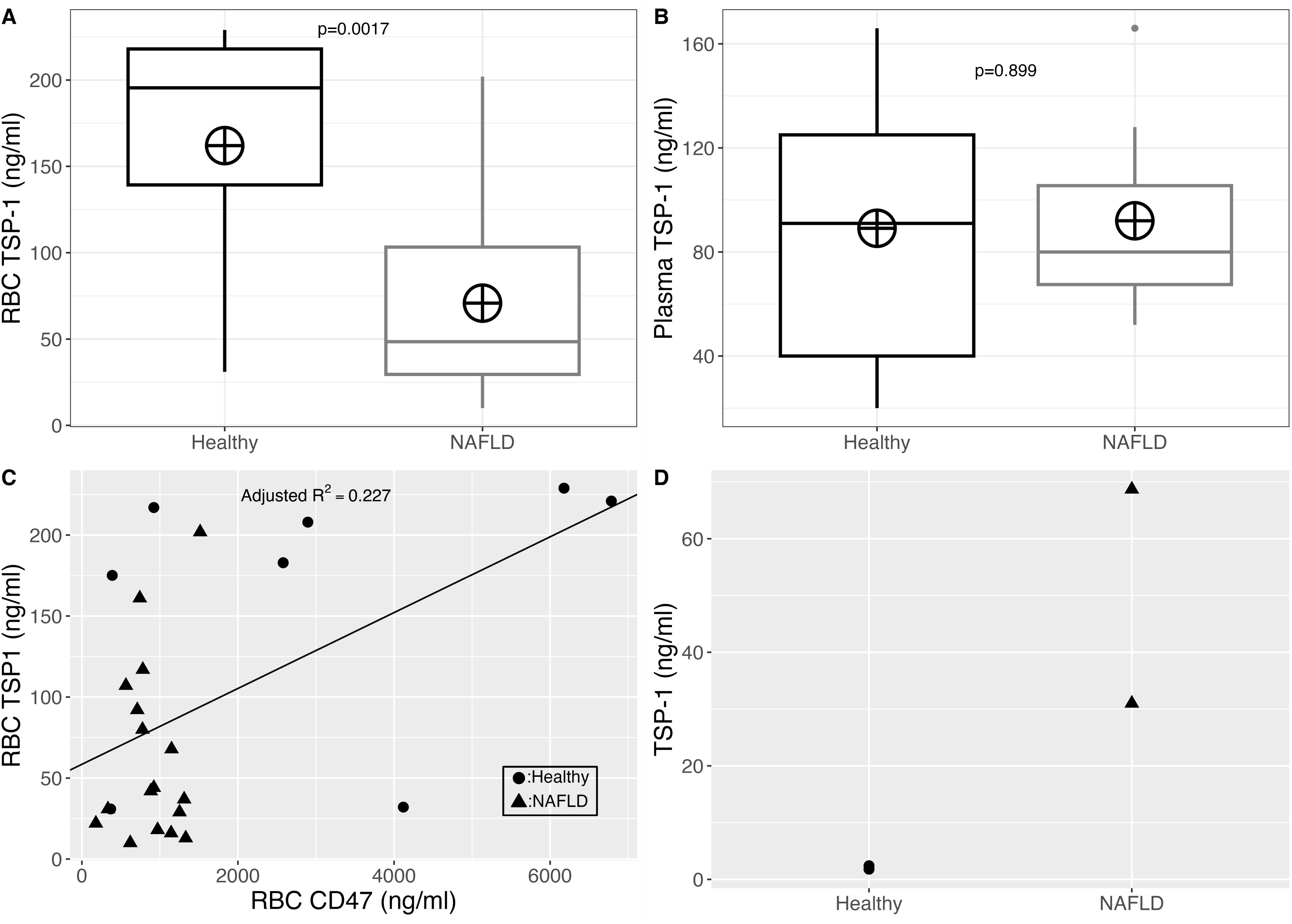
A. Erythrocyte TSP-1 levels in in healthy controls (n=8) and MAFLD patients (n=22). B. Erythrocyte TSP-1 levels in in healthy controls (n=9) and MAFLD patients (n=7). C. Linear regression between erythrocyte CD47 and erythrocyte TSP-1 levels. D. TSP-1 levels in the conditioned media of erythrocytes of healthy controls (n=2) and MAFLD patients (n=2) incubated with 80 ng/ml of recombinant TSP-1. CD47: cluster of differentiation 47; MAFLD: metabolic associated fatty liver disease; RBC: red blood cells; TPS-1: thrombospondin-1.

### INCUBATION OF ERYTHROCYTES WITH TSP-1

In an in vitro experiment, we incubated erythrocytes of 2 healthy controls or 2 MAFLD patients with recombinant TSP-1. We showed that erythrocyte-conditioned media of MAFLD patients contained higher TSP-1 levels, indicating lower TSP-1 binding (figure 3). This result agrees with our previous results of lower TSP-1 on erythrocytes of MAFLD patients.

### ERYTHROCYTE ARGINASE-1 PROTEIN AND ACTIVITY LEVELS

CD47 ligation by TSP-1 impairs the production of nitric oxide^25^. Since the levels of TSP-1 were lower in erythrocytes of MAFLD patients, we speculated that the synthesis of nitric oxide could be downregulated. Because arginase-1 is an important modulator of the nitric oxide levels on erythrocytes, we wondered whether its protein levels are reduced in erythrocytes of MAFLD patients. As shown in Table 2, and Figure4 MAFLD patients had lower erythrocyte arginase-1 levels (mean=156 ng/ml) than healthy subjects (mean=170 ng/ml), with a statistically significant difference (p=0.0092) and a large effect size but low statistical power 1-β=0.21. Similar results were obtained for the enzymatic activity of arginase-1 (Table 2, Figure4). ARG-1 enzymatic activity was lower in MAFLD patients (mean=7.0 U/ml), compared to healthy subjects (mean=7.7 U/ml). The difference was statistically significant with p=0.042.

In addition, the 95%CI for the difference between the means does not contain the zero value. This means that although the statistical power is low, we can, however, be confident that there is a statistically significant difference.

Thus, we hypothesize that, erythrocyte arginase expression is downregulated in MAFLD and this contradicts previous reports with regards to diabetes melitus^35,36^.

### PLASMA ARGINASE-1 LEVELS

Erythrocytes can take-up arginine from their environment. Thus, the levels of available arginine for the production of erythrocytic nitric oxide can be also regulated by the levels of plasma arginase. As such, we determined the protein levels of arginase-1 in the plasma of MAFLD patients and healthy controls. There was no statistically significant difference in plasma arginase-1 between MAFLD patients (mean=65 ng/ml) and healthy subjects (mean=52 ng/ml), with p=0.691 (Table 2, and Figure 4).

**Figure 4:**
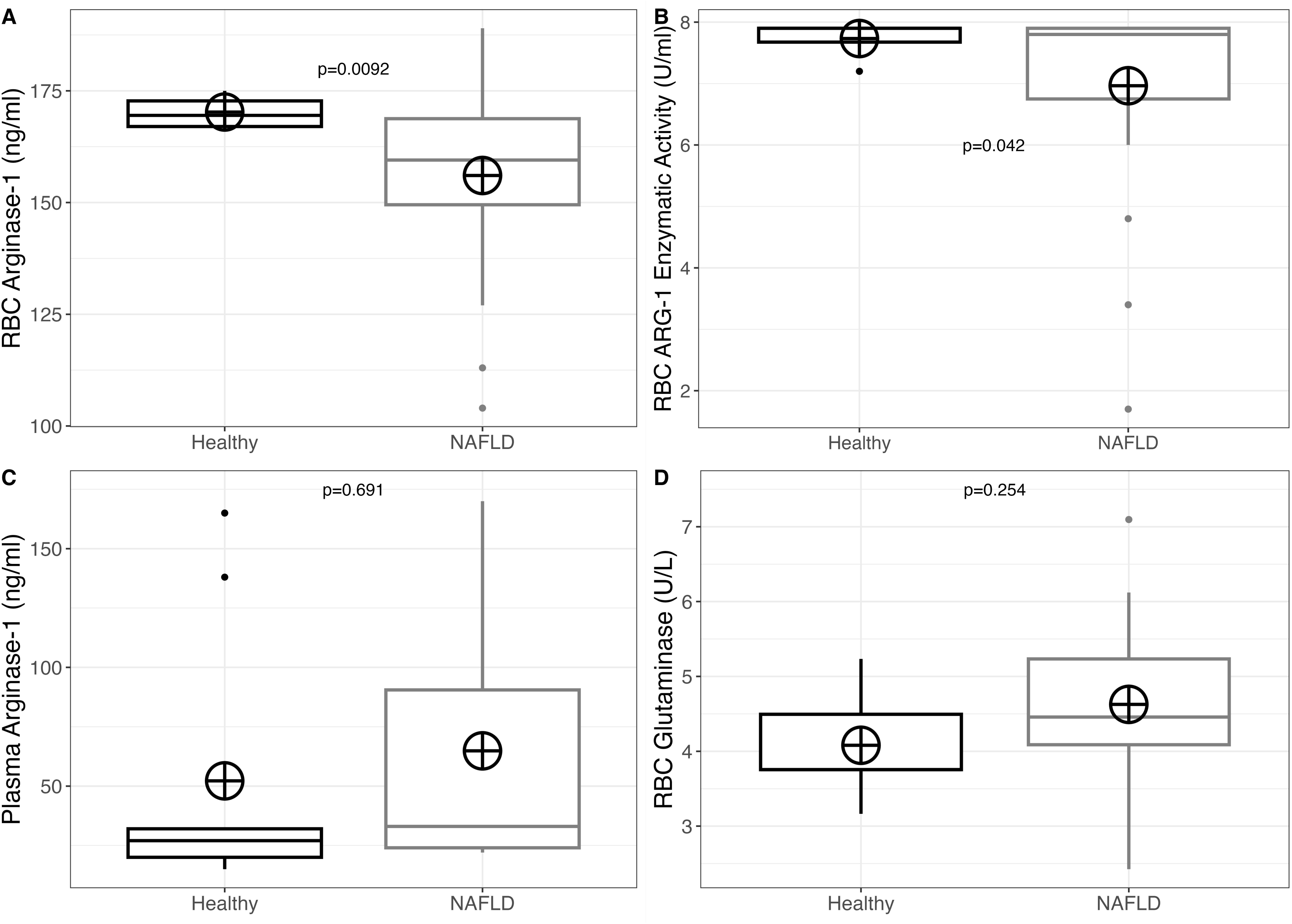
Erythrocyte ARG-1 protein levels in healthy controls (n=4) and MAFLD patients (n=22). B. Erythrocyte ARG-1 activity levels in healthy controls (n=6) and MAFLD patients (n=23). C. Plasma ARG-1 protein levels in healthy controls (n=9) and MAFLD patients (n=7). D. Erythrocyte glutaminase activity in erythrocytes of healthy controls (n=5) and MAFLD patients (n=18). ARG-1: arginase-1, RBC: red blood cells; MAFLD: metabolic associated fatty liver disease.

### ERYTHROCYTE GLUTAMINASE ACTIVITY

Erythrocytes contain a bioactive glutaminase. Its role is mainly indicated in the synthesis of glutamate for the de novo synthesis of glutathione and the transamination of pyruvate^37^. In MAFLD, have previously been reported increased hepatic glutaminase activity and increased serum glutamate to glutamine ratio^38^. Based on these, we speculated that erythrocyte glutaminase could contribute to the overall increased glutaminolysis in MAFLD. We found that erythrocyte glutaminase is increased in MAFLD patients, albeit the difference is not statistically significant (p=0.25) (table 2 and figure 4).

## DISCUSSION

MAFLD constitutes a metabolic hepatic disease, where metabolic perturbations precede inflammatory mechanisms. Otogawa et al^18^ and Park et al^19^, have reported that hepatic erythrophagocytosis results in inflammation during MAFLD. We have also reported extended erythrocyte changes during MAFLD^20–22,39^. In this study we try to elaborate on the role of erythrocytes in the metabolic inflammation of MAFLD.

Lipotoxicity induces TLR9 upregulation^6,7^, and activation in the liver through hepatocyte mitochondrial DNA release^7,9^,. This step actually connects dysmetabolism with metabolic inflammation in the liver during MAFLD^37^. Similarly, reduced TLR9 levels on T-lymphocytes protect from the transition from hepatic steatosis to metabolic inflammation^13^. Of note, a previous study reported that, in general, mitochondrial DNA-induced inflammation requires erythrocyte TLR9, and erythrophagocytosis^9^. In this study, we show that erythrocytes of MAFLD patients contain increased TLR9 levels and follow the trend of upregulated TLR9 in the liver of MAFLD animals and patients^6,7^. In this perspective, erythrocytes join hepatocytes, liver macrophages^9^, endothelial cells^40^, T-cells^13^ and hepatic stellate cells^12^ in sensing liver damage through TLR9. Erythrocytes with high TLR9 levels could bind the increased cell-free mitochondrial DNA^8,9^, and deliver it to the liver through their phagocytosis, which is augmented via CD47 loss^21^, phosphatidylserine exposure^18^, MCP1 release^22^ and sphingosine accumulation^21^. Accordingly, increased mitochondrial DNA delivery through erythrocyte TLR9 in MAFLD, could lead to the formation of a specific pro-inflammatory erythrophagocyte-macrophage subset, though chronic macrophage TLR9 stimulation^41^.

Our hypothesis of TLR9-induced pro-inflammatory erythrophagocytosis seems to be contradictory to previous studies with regard to MAFLD. In particular, previous studies claimed that solely iron is responsible for hepatic erythrophagocytosis-induced inflammation^18,19^. However, iron alone is not sufficient for triggering ferroptosis and inflammation. Recent reports indicated that erythrocyte lipid peroxides, sphingosine and erythrocyte-bound mitochondrial DNA render erythrophagocytosis pro-inflammatory^23,42,43^. In the past, we have shown that erythrocytes of MAFLD patients contain increased sphingosine^21^. In this study, we report increased TLR9 levels on erythrocytes, which could contribute to the increased delivery of mitochondrial DNA to macrophages, resulting in inflammation^23^. However, we should keep in mind that systemic metabolic perturbation could amplify erythrocyte-dependent inflammation irrespectively of TLR9^44–46^. Erythrocyte TLR9 upregulation probably stems from the effort of the organism to scavenge the circulating cell-free mitochondrial DNA^47^.

We also show that TLR9 levels are associated with erythrocyte sphingosine levels. One way to interpret this correlation is that sphingosine leads to TLR9 accumulation in erythrocytes of MAFLD patients. Relatively, a recent study found that cholesterol impairs TLR9 degradation^15^. Mechanistically, this is preceded by sphingosine accumulation-induced invaginations and endolysosome dysfunction^14^. Accordingly, sphingolipid accumulation has been shown to be necessary for erythrocyte invaginations^48,49^; erythrocyte invaginations are enriched in BAND3, a protein that exhibits physical contact with TLR9 in erythrocytes^23^. We hypothesize that sphingosine could partially explain TLR9 accumulation through TLR9 entrapment in invaginations or endosomes. Alternatively, TLR9 activation could lead to sphingosine formation as previously described in lung epithelial cells^50^.

Our result of upregulated erythrocyte TLR9, along with the increased mitochondrial DNA in the sera of MAFLD patients as previously reported^8,9^ led us to hypothesize that MAFLD erythrocytes would also exhibit increased TLR9 activation. Thus, we focused on the effects that this event could have.

A previous report showed that part of the effect of erythrocyte TLR9 activation can be recapitulated by lipid remodeling^23^, through hydrogen peroxide which reduces the PC/PE ratio^51^. Hence, we explored the PC/PE ratio in erythrocyte of MAFLD patients. We show that this ratio decreases in erythrocytes of MAFLD patients, agreeing with a previous report^52^. Importantly, this ratio negatively correlated with TLR9 levels.

We speculate that the PC/PE ratio correlates with TLR9 levels because increased erythrocyte TLR9 levels would sense the increased cell-free mitochondrial DNA in MAFLD^6,7^ and would lead to lipid remodeling. However, since erythrocytes also reflect the systemic lipotoxicity^45^, we hypothesize that, in part, the reduced PC/PE ratio in erythrocytes is caused by overall lipid changes in the body. Yet, one could argue that the reduced erythrocyte PC/PE ratio is caused by systemic inflammation-mediated selective PC hydrolysis, as previously shown^53^. However, our results of increased PE content in erythrocyte membranes of MAFLD patients partially exclude this mechanism.

Independently of the cause, reduced erythrocyte PC/PE ratio could participate in the lipotoxic signals after erythrophagocytosis. In particular, reduced PC/PE ratio has been implicated in the transition of steatosis to steatohepatitis, since it is linked to reduced triglyceride export from the liver, loss of hepatic structural integrity, endoplasmic reticulum stress, dysregulated mitochondrial function and inflammation^54–59^. Furthermore, phosphatidylethanolamine is linked to steatosis, dysregulated autophagy, apoptosis, oxidative stress and inflammation^60^.

Previous reports have proved that erythrocyte TLR9 activation leads to CD47 loss^23^. Relatively, we have reported in the past decreased CD47 in erythrocytes of MAFLD patients^21^. As expected, TLR9 levels were inversely correlated with CD47 levels. We excluded that aging causes CD47 loss, by measuring the levels of CD35, a molecule whose levels decline with erythrocyte aging^33^. This result of unaltered CD35 levels also excludes the contribution of erythrocyte to complement-induced tissue damage, in contrast to hepatitis C^61^. We hypothesize, thus, that TLR9 upregulation and activation would lead to greater CD47 loss. This could also favor hepatic erythrophagocytosis^62^, a major mechanism driving inflammation in MAFLD^18^. Based on these, we propose that TLR9-induced CD47 loss could also favor the delivery of the erythrocyte aberrant lipidome to macrophages. In addition, TLR9-induced erythrocyte CD47 loss could partially explain why anti-CD47 treatment in MAFLD leads to anemia^63^; the already reduced erythrocyte CD47 levels would be blocked leading to extended erythrophagocytosis.

Erythrocyte surface protein loss is accompanied by chemokine release^24^. Based on the correlation of TLR9 and CD47, we hypothesized that TLR9 would also correlate positively with the release of MCP-1. In a previous study we reported increased MCP-1 release from MAFLD erythrocytes^22^. Here, we found that, indeed, the more the TLR9 the more the MCP-1 release. This result is of major importance since MCP-1 participates in the transition from hepatic steatosis to steatohepatitis^64^, and implies the existence of a proinflammatory TLR9-MCP-1 pathway in erythrocytes previously reported for other cell types^65,66^.

Erythrocyte CD47 binds TSP-1^34^, producing a pro-phagocytosis signal^67^. Thus, we hypothesized that the reduced CD47 levels could affect the TSP-1 binding. We show that TSP-1 levels were reduced in erythrocyte of MAFLD patients. In addition, we excluded the scenario that the reduced levels on erythrocytes are caused by reduced circulating TSP-1 levels in the plasma. The levels of TSP-1 were positively correlated with CD47 and inversely correlated with TLR9 levels. In vitro, erythrocytes of MAFLD patients bound less TSP-1. We speculate that reduced TSP-1 binding would allow more TSP-1 molecules to remain in the liver and drive metabolic inflammation^68^. In contrast to our results, previous studies have connected erythrocyte CD47 loss to increased TSP-1 binding^34^. However, our results should not be considered conflicting. CD47 loss is not always accompanied by increased TSP-1 binding^34^. Thus, in MAFLD, probably CD47 determines the degree of TSP-1 binding. Our report of reduced TSP-1 levels on erythrocytes, suggests that TSP-1 does not drive hepatic erythrophagocytosis in MAFLD. Nevertheless, we should keep in mind that TSP-1 also binds to CD36^69^. Hence, its levels could also influence TSP-1 binding.

Hepatic lipotoxicity is supported by amino acid metabolism deregulation^4,5^. For this reason, we also focused on the metabolism of arginine and glutamine in erythrocytes during MAFLD. We show that the reduced CD47-TSP-1 levels are accompanied by reduced arginase-1 protein and activity levels. This is no surprise since it has been shown in the past that CD47 ligation by TSP-1 negatively regulates nitric oxide synthesis^25^. However, this contradicts the results with regard to erythrocyte arginase-1 in diabetes^35,36^. Mechanistically, reduced arginase-1 activity could increase the availability of arginine for the synthesis of nitric oxide. Importantly, decreased arginase activity drives the impairment of intestinal barrier during MAFLD. Remarkably, this event is associated with beginning hepatic inflammation.^5^ This mechanism of reduced CD47-TSP-1 binding-driven metabolic inflammation, partially explains the effect of long-term CD47 deficiency on steatohepatitis^70^.

We next focused on glutaminase metabolism. In MAFLD, there is a marked increased glutamate to glutamine ratio in the sera of MAFLD patients^38^. In the liver, increased glutaminase drives the formation of an increased a-ketoglutarate pool; this leads to MAFLD through reduction of the PC/PE ratio^4^. Relatively, TSP-1 negatively impacts the metabolism of glutamine^26^. In erythrocytes, glutamine is used for de novo synthesis of the antioxidant glutathione and transamination of pyruvate^37^. As such, a previous study found that erythrocyte TSP-1 binding led to increased reactive oxygen species concentration^25^. Based on these, we speculated that reduce d TSP-1 levels on the erythrocytes of MAFLD patients would be correlated with increased erythrocyte glutaminase. We found increased glutaminase activity on the erythrocytes of MAFLD patients, but this difference was not statistically significant. Based on our calculations, a higher number of participants is needed for the difference to appear significant.

Why would these mechanisms be evolved?These mechanisms would lead to mitochondrial DNA scavenging (through TLR9 upregulation) and elimination (through induction of erythrophagocytosis) inhibiting immune cell activation. However, the extended activation under chronic inflammation exacerbates tissue damage.

Our study has some limitations. Firstly, the number of the participants is low. Nevertheless, we have used robust mathematical models to overcome this downside. Secondly, we did not discriminate between liver steatosis and steatohepatitis though liver biopsy. However, all patients had steatosis and elevated liver enzymes, signifying liver damage. In addition, we did not quantify the levels of erythrocyte-bound mitochondrial DNA. Nevertheless, it is rational to hypothesize that increased erythrocyte TLR9 levels along with increased circulating cell-free mitochondrial DNA levels in MAFLD^8^, can result in augmented erythrocyte-bound mtDNA levels. Furthermore, we did not discriminate between intracellular and surface TLR9 localization. We should note though, that both surface and intracellular erythrocyte TLR9 can bind mitochondrial DNA^47^; a proportion of TLR9 molecules of erythrocyte must be on the surface^47^. Finally, we do not provide a mechanistic insight for the correlations found. Despite the accumulated bibliography supporting our interpretations, it is possible that these correlations are the outcome of inflammatory mechanisms acting simultaneously.

Overall, we report here that during MAFLD, TLR9, is upregulated in erythrocytes and is associated with sphingosine, and reduced erythrocyte PC/PE ratio. Increased erythrocyte TLR9 levels were followed by decreased CD47, increased MCP1 release and reduced scavenging of TSP-1. These changes are followed by decreased erythrocyte arginase-1 and a minor increase of erythrocyte glutaminase-1 activity.

## CONCLUSIONS

We propose that in MAFLD, erythrocyte TLR9 upregulation mediates metabolic inflammation, through its associations with erythrocyte sphingosine and glycerophospholipid perturbation, CD47 loss, MCP1 release, reduced TSP-1 scavenging, decreased arginase and increased glutaminase (figure 5).

**Figure 5:**
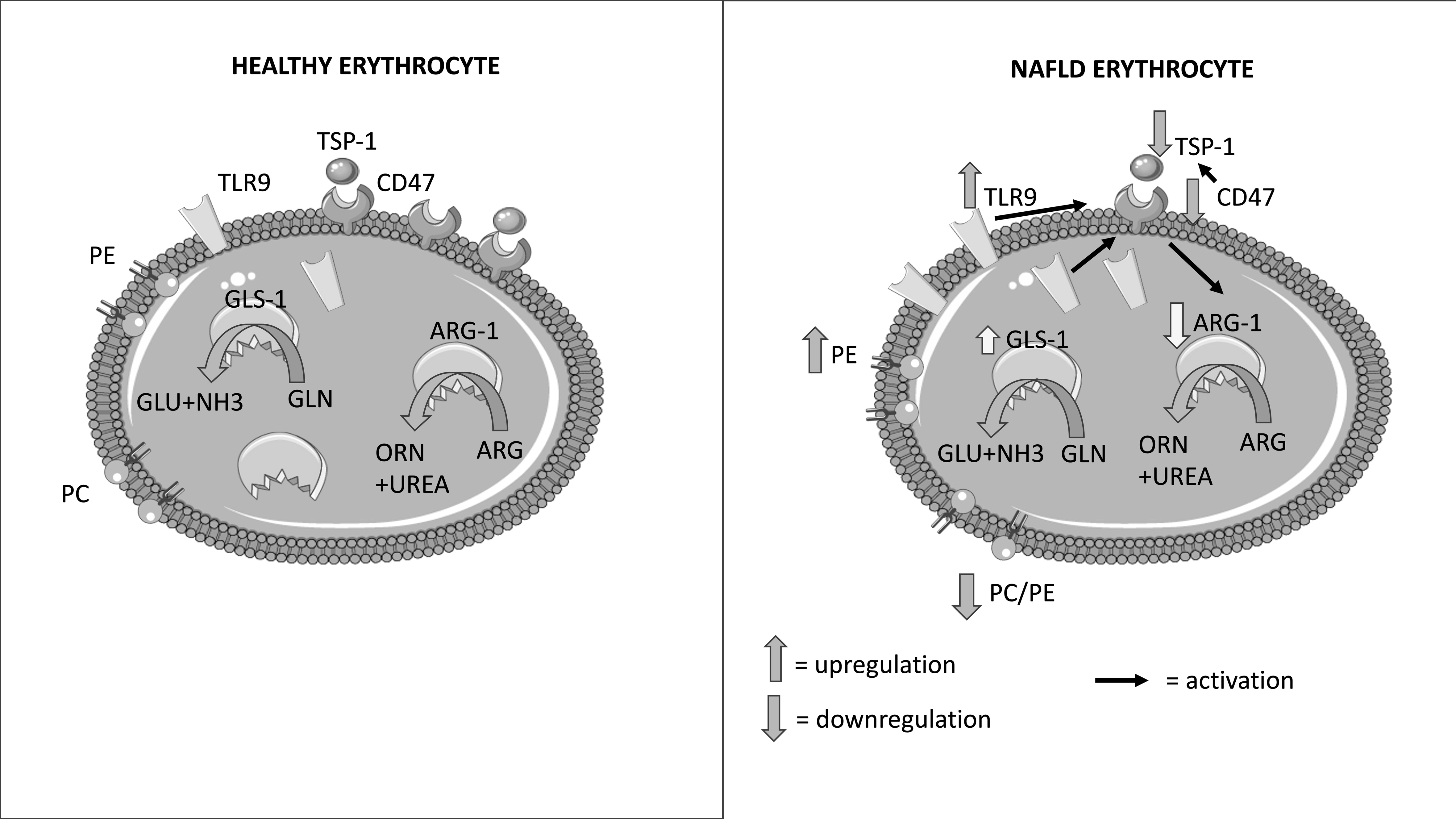
In MAFLD, TLR9 is upregulated in erythrocytes and is associated with sphingosine, and reduced erythrocyte PC/PE ratio. Increased erythrocyte TLR9 levels are followed by decreased CD47, increased MCP1 release and reduced scavenging of TSP-1. These changes are followed by decreased erythrocyte arginase-1 and a minor increase of erythrocyte glutaminase-1 activity. ARG-1: arginase-1; GLS-1: glutaminase-1; MAFLD: metabolic dysfunction fatty liver disease; PC: phosphatidylcholine; PE: phosphatidylethanolamine; RBC: red blood cells; TPS-1: thrombospondin-1; TLR9: toll-like receptor 9.

**Author Contributions:** Conceptualization, C.P; methodology, C.P; software, K.A; validation, C.P, KA.; formal analysis, C.P, K.A; investigation, C.P.; resources K.M; K.A; data curation, C.P, K.A.; writing—original draft preparation, C.P.; writing—review and editing, C.P, K.A, K.M; supervision, C.P, K.A.; project administration, C.P, K.A, I.T, K.M; funding acquisition, C.P, K.A, K.M. All authors have read and agreed to the published version of the manuscript.

**Institutional Review Board Statement:** The study was conducted in accordance with the Declaration of Helsinki, and approved by the Ethics Committee of University Hospital of Alexandroupolis (72/23-01-2018 l 02-04-2018).

**Informed Consent Statement:** Informed consent was obtained from all subjects involved in the study.

**Conflicts of Interest:** The authors declare no conflict of interest

**Funding:** This work is funded by the Hellenic Association for the Study of the Liver

## ABBREVIATIONS

ALT: alanine transaminase
ARG-1: arginase-1
AST: aspartate transaminase
CD47: cluster of differentiation 47
CHOL: cholesterol
MAFLD: metabolic dysfunction associated fatty liver disease
MCP-1: monocyte chemoattractant protein 1
PC: phosphatidylcholine
PE: phosphatidylethanolamine
TLR9: toll-like receptor 9
9TSP-1: thombospondin-1

---

Figure S1: Linear regression between erythrocyte TLR9 and erythrocyte sphingosine levels. Sph: sphingosine; TLR9: toll-like receptor 9.

## REFERENCES

(1) Blüher, M. Obesity: Global Epidemiology and Pathogenesis. Nat. Rev. Endocrinol. 2019, 15 (5), 288–298. 10.1038/s41574-019-0176-8.

(2) Papadopoulos, C.; Tentes, I.; Papazoglou, D.; Anagnostopoulos, K. Lysophospholipid Metabolism and Signalling in Non-Alcoholic Fatty Liver Disease. Folia Med. (Plovdiv) 2022, 64 (1), 7–12. 10.3897/folmed.64.e59297.

(3) Caligiuri, A.; Gentilini, A.; Marra, F. Molecular Pathogenesis of NASH. Int. J. Mol. Sci. 2016, 17 (9), 1575. 10.3390/ijms17091575.

(4) Simon, J.; Nuñez-García, M.; Fernández-Tussy, P.; Barbier-Torres, L.; Fernández-Ramos, D.; Gómez-Santos, B.; Buqué, X.; Lopitz-Otsoa, F.; Goikoetxea-Usandizaga, N.; Serrano-Macia, M.; Rodriguez-Agudo, R.; Bizkarguenaga, M.; Zubiete-Franco, I.; Juan, V. G.; Cabrera, D.; Alonso, C.; Iruzubieta, P.; Romero-Gomez, M.; van Liempd, S.; Castro, A.; Nogueiras, R.; Varela-Rey, M.; Falcón-Pérez, J. M.; Villa, E.; Crespo, J.; Lu, S. C.; Mato, J. M.; Aspichueta, P.; Delgado, T. C.; Martínez-Chantar, M. L. Targeting Hepatic Glutaminase 1 Ameliorates Non-Alcoholic Steatohepatitis by Restoring Very-Low Density Lipoproteins Triglyceride Assembly. Cell Metab. 2020, 31 (3), 605–622.e10. 10.1016/j.cmet.2020.01.013.

(5) Baumann, A.; Rajcic, D.; Brandt, A.; Sánchez, V.; Jung, F.; Staltner, R.; Nier, A.; Trauner, M.; Staufer, K.; Bergheim, I. Alterations of Nitric Oxide Homeostasis as Trigger of Intestinal Barrier Dysfunction in Non-Alcoholic Fatty Liver Disease. J. Cell. Mol. Med. 2022, 26 (4), 1206–1218. 10.1111/jcmm.17175.

(6) Sawada, K.; Ohtake, T.; Hasebe, T.; Abe, M.; Tanaka, H.; Ikuta, K.; Suzuki, Y.; Fujiya, M.; Hasebe, C.; Kohgo, Y. Augmented Hepatic Toll-like Receptors by Fatty Acids Trigger the pro-Inflammatory State of Non-Alcoholic Fatty Liver Disease in Mice. Hepatol. Res. Off. J. Jpn. Soc. Hepatol. 2014, 44 (8), 920–934. 10.1111/hepr.12199.

(7) Mridha, A. R.; Haczeyni, F.; Yeh, M. M.; Haigh, W. G.; Ioannou, G. N.; Barn, V.; Ajamieh, H.; Adams, L.; Hamdorf, J. M.; Teoh, N. C.; Farrell, G. C. TLR9 Is Up-Regulated in Human and Murine NASH: Pivotal Role in Inflammatory Recruitment and Cell Survival. Clin. Sci. Lond. Engl. 1979 2017, 131 (16), 2145–2159. 10.1042/CS20160838.

(8) Gao, Y.; Wang, Y.; Liu, H.; Liu, Z.; Zhao, J. Mitochondrial DNA from Hepatocytes Induces Upregulation of Interleukin-33 Expression of Macrophages in Nonalcoholic Steatohepatitis. Dig. Liver Dis. Off. J. Ital. Soc. Gastroenterol. Ital. Assoc. Study Liver 2020, 52 (6), 637–643. 10.1016/j.dld.2020.03.021.

(9) Garcia-Martinez, I.; Santoro, N.; Chen, Y.; Hoque, R.; Ouyang, X.; Caprio, S.; Shlomchik, M. J.; Coffman, R. L.; Candia, A.; Mehal, W. Z. Hepatocyte Mitochondrial DNA Drives Nonalcoholic Steatohepatitis by Activation of TLR9. J. Clin. Invest. 2016, 126 (3), 859– 864. 10.1172/JCI83885.

(10) Miura, K.; Kodama, Y.; Inokuchi, S.; Schnabl, B.; Aoyama, T.; Ohnishi, H.; Olefsky, J. M.; Brenner, D. A.; Seki, E. Toll-like Receptor 9 Promotes Steatohepatitis by Induction of Interleukin-1beta in Mice. Gastroenterology 2010, 139 (1), 323–334.e7. 10.1053/j.gastro.2010.03.052.

(11) An, P.; Wei, L.-L.; Zhao, S.; Sverdlov, D. Y.; Vaid, K. A.; Miyamoto, M.; Kuramitsu, K.; Lai, M.; Popov, Y. V. Hepatocyte Mitochondria-Derived Danger Signals Directly Activate Hepatic Stellate Cells and Drive Progression of Liver Fibrosis. Nat. Commun. 2020, 11 (1), 2362. 10.1038/s41467-020-16092-0.

(12) Watanabe, A.; Hashmi, A.; Gomes, D. A.; Town, T.; Badou, A.; Flavell, R. A.; Mehal, W. Z. Apoptotic Hepatocyte DNA Inhibits Hepatic Stellate Cell Chemotaxis via Toll-like Receptor 9. Hepatol. Baltim. Md 2007, 46 (5), 1509–1518. 10.1002/hep.21867.

(13) Alegre, N. S.; Garcia, C. C.; Billordo, L. A.; Ameigeiras, B.; Poncino, D.; Benavides, J.; Colombato, L.; Cherñavsky, A. C. Limited Expression of TLR9 on T Cells and Its Functional Consequences in Patients with Nonalcoholic Fatty Liver Disease. Clin. Mol. Hepatol. 2020, 26 (2), 216–226. 10.3350/cmh.2019.0074.

(14) Lima, S.; Milstien, S.; Spiegel, S. Sphingosine and Sphingosine Kinase 1 Involvement in Endocytic Membrane Trafficking. J. Biol. Chem. 2017, 292 (8), 3074–3088. 10.1074/jbc.M116.762377.

(15) Teratani, T.; Tomita, K.; Suzuki, T.; Furuhashi, H.; Irie, R.; Hida, S.; Okada, Y.; Kurihara, C.; Ebinuma, H.; Nakamoto, N.; Saito, H.; Hibi, T.; Miura, S.; Hokari, R.; Kanai, T. Free Cholesterol Accumulation in Liver Sinusoidal Endothelial Cells Exacerbates Acetaminophen Hepatotoxicity via TLR9 Signaling. J. Hepatol. 2017, 67 (4), 780–790. 10.1016/j.jhep.2017.05.020.

(16) Papadopoulos, C.; Panopoulou, M.; Anagnostopoulos, K.; Tentes, I. Immune and Metabolic Interactions of Human Erythrocytes: A Molecular Perspective. Endocr. Metab. Immune Disord. Drug Targets 2021, 21 (5), 843–853. 10.2174/1871530320666201104115016.

(17) Papadopoulos, C.; Tentes, I.; Anagnostopoulos, K. Erythrocytes Contribute to the Immunometabolic Cross-Talk. Immunometabolism 2021, 3 (2), e210015. 10.20900/immunometab20210015.

(18) Otogawa, K.; Kinoshita, K.; Fujii, H.; Sakabe, M.; Shiga, R.; Nakatani, K.; Ikeda, K.; Nakajima, Y.; Ikura, Y.; Ueda, M.; Arakawa, T.; Hato, F.; Kawada, N. Erythrophagocytosis by Liver Macrophages (Kupffer Cells) Promotes Oxidative Stress, Inflammation, and Fibrosis in a Rabbit Model of Steatohepatitis: Implications for the Pathogenesis of Human Nonalcoholic Steatohepatitis. Am. J. Pathol. 2007, 170 (3), 967–980. 10.2353/ajpath.2007.060441.

(19) Park, J. B.; Ko, K.; Baek, Y. H.; Kwon, W. Y.; Suh, S.; Han, S.-H.; Kim, Y. H.; Kim, H. Y.; Yoo, Y. H. Pharmacological Prevention of Ectopic Erythrophagocytosis by Cilostazol Mitigates Ferroptosis in NASH. Int. J. Mol. Sci. 2023, 24 (16), 12862. 10.3390/ijms241612862.

(20) Papadopoulos, C.; Mimidis, K.; Tentes, I.; Tente, T.; Anagnostopoulos, K. Validation and Application of a Protocol for the Extraction and Quantitative Analysis of Sphingomyelin in Erythrocyte Membranes of Patients with Non-Alcoholic Fatty Liver Disease. JPC – J. Planar Chromatogr. – Mod. TLC 2021, 34 (5), 411–418. 10.1007/s00764-021-00127-3.

(21) Papadopoulos, C.; Spourita, E.; Mimidis, K.; Kolios, G.; Tentes, L.; Anagnostopoulos, K. Nonalcoholic Fatty Liver Disease Patients Exhibit Reduced CD47 and Increased Sphingosine, Cholesterol, and Monocyte Chemoattractant Protein-1 Levels in the Erythrocyte Membranes. Metab. Syndr. Relat. Disord. 2022, 20 (7), 377–383. 10.1089/met.2022.0006.

(22) Papadopoulos, C.; Mimidis, K.; Papazoglou, D.; Kolios, G.; Tentes, I.; Anagnostopoulos, K. Red Blood Cell-Conditioned Media from Non-Alcoholic Fatty Liver Disease Patients Contain Increased MCP1 and Induce TNF-α Release. Rep. Biochem. Mol. Biol. 2022, 11 (1), 54–62. 10.52547/rbmb.11.1.54.

(23) Lam, L. K. M.; Murphy, S.; Kokkinaki, D.; Venosa, A.; Sherrill-Mix, S.; Casu, C.; Rivella, S.; Weiner, A.; Park, J.; Shin, S.; Vaughan, A.; Hahn, B. H.; Odom John, A. R.; Meyer, N. J.; Hunter, C. A.; Worthen, G. S.; Mangalmurti, N. S. DNA Binding to TLR9 Expressed by Red Blood Cells Promotes Innate Immune Activation and Anemia. Sci. Transl. Med. 2021, 13 (616), eabj1008. 10.1126/scitranslmed.abj1008.

(24) Dumaswala, U. J.; Zhuo, L.; Mahajan, S.; Nair, P. N.; Shertzer, H. G.; Dibello, P.; Jacobsen, D. W. Glutathione Protects Chemokine-Scavenging and Antioxidative Defense Functions in Human RBCs. Am. J. Physiol. Cell Physiol. 2001, 280 (4), C867–873. 10.1152/ajpcell.2001.280.4.C867.

(25) Bissinger, R.; Petkova-Kirova, P.; Mykhailova, O.; Oldenborg, P.-A.; Novikova, E.; Donkor, D. A.; Dietz, T.; Bhuyan, A. A. M.; Sheffield, W. P.; Grau, M.; Artunc, F.; Kaestner, L.; Acker, J. P.; Qadri, S. M. Thrombospondin-1/CD47 Signaling Modulates Transmembrane Cation Conductance, Survival, and Deformability of Human Red Blood Cells. Cell Commun. Signal. CCS 2020, 18 (1), 155. 10.1186/s12964-020-00651-5.

(26) Bronson, S. M.; Westwood, B.; Cook, K. L.; Emenaker, N. J.; Chappell, M. C.; Roberts, D. D.; Soto-Pantoja, D. R. Discrete Correlation Summation Clustering Reveals Differential Regulation of Liver Metabolism by Thrombospondin-1 in Low-Fat and High-Fat Diet-Fed Mice. Metabolites 2022, 12 (11), 1036. 10.3390/metabo12111036.

(27) Gwag, T.; Reddy Mooli, R. G.; Li, D.; Lee, S.; Lee, E. Y.; Wang, S. Macrophage-Derived Thrombospondin 1 Promotes Obesity-Associated Non-Alcoholic Fatty Liver Disease. JHEP Rep. Innov. Hepatol. 2021, 3 (1), 100193. 10.1016/j.jhepr.2020.100193.

(28) Folch, J.; Lees, M.; Sloane Stanley, G. H. A Simple Method for the Isolation and Purification of Total Lipides from Animal Tissues. J. Biol. Chem. 1957, 226 (1), 497–509.

(29) *[PDF] R: A language and environment for statistical computing. | Semantic Scholar*. https://www.semanticscholar.org/paper/R%3A-A-language-and-environment-for-statistical-Team/659408b243cec55de8d0a3bc51b81173007aa89b (accessed 2023-11-25).

(30) Welch, B. The Generalisation of Student’s Problems When Several Different Population Variances Are Involved. Biometrika 1947, No. 34, 28–35.

(31) Gignac, G. E.; Szodorai, E. T. Effect Size Guidelines for Individual Differences Researchers. Personal. Individ. Differ. 2016, No. 102, 74–78.

(32) Eldesouky, N. A.; Abo El Fetouh, R. M.; Hafez, A. A.; Gad, A.; Kamal, M. M. The Expression of CD47 and Its Association with 2,3-DPG Levels in Stored Leuco-Reduced Blood Units. Transfus. Clin. Biol. J. Soc. Francaise Transfus. Sang. 2019, 26 (4), 279–283. 10.1016/j.tracli.2019.01.004.

(33) Ripoche, J.; Sim, R. B. Loss of Complement Receptor Type 1 (CR1) on Ageing of Erythrocytes. Studies of Proteolytic Release of the Receptor. Biochem. J. 1986, 235 (3), 815–821. 10.1042/bj2350815.

(34) Wang, F.; Liu, Y.-H.; Zhang, T.; Gao, J.; Xu, Y.; Xie, G.-Y.; Zhao, W.-J.; Wang, H.; Yang, Y.-G. Aging-Associated Changes in CD47 Arrangement and Interaction with Thrombospondin-1 on Red Blood Cells Visualized by Super-Resolution Imaging. Aging Cell 2020, 19 (10), e13224. 10.1111/acel.13224.

(35) Zhou, Z.; Mahdi, A.; Tratsiakovich, Y.; Zahorán, S.; Kövamees, O.; Nordin, F.; Uribe Gonzalez, A. E.; Alvarsson, M.; Östenson, C.-G.; Andersson, D. C.; Hedin, U.; Hermesz, E.; Lundberg, J. O.; Yang, J.; Pernow, J. Erythrocytes From Patients With Type 2 Diabetes Induce Endothelial Dysfunction Via Arginase I. J. Am. Coll. Cardiol. 2018, 72 (7), 769– 780. 10.1016/j.jacc.2018.05.052.

(36) Mahdi, A.; Tengbom, J.; Alvarsson, M.; Wernly, B.; Zhou, Z.; Pernow, J. Red Blood Cell Peroxynitrite Causes Endothelial Dysfunction in Type 2 Diabetes Mellitus via Arginase. Cells 2020, 9 (7), 1712. 10.3390/cells9071712.

(37) Reisz, J. A.; Slaughter, A. L.; Culp-Hill, R.; Moore, E. E.; Silliman, C. C.; Fragoso, M.; Peltz, E. D.; Hansen, K. C.; Banerjee, A.; D’Alessandro, A. Red Blood Cells in Hemorrhagic Shock: A Critical Role for Glutaminolysis in Fueling Alanine Transamination in Rats. Blood Adv. 2017, 1 (17), 1296–1305. 10.1182/bloodadvances.2017007187.

(38) Du, K.; Chitneni, S. K.; Suzuki, A.; Wang, Y.; Henao, R.; Hyun, J.; Premont, R. T.; Naggie, S.; Moylan, C. A.; Bashir, M. R.; Abdelmalek, M. F.; Diehl, A. M. Increased Glutaminolysis Marks Active Scarring in Nonalcoholic Steatohepatitis Progression. Cell. Mol. Gastroenterol. Hepatol. 2019, 10 (1), 1–21. 10.1016/j.jcmgh.2019.12.006.

(39) Papadopoulos, C.; Tentes, I.; Anagnostopoulos, K. Red Blood Cell Dysfunction in Non-Alcoholic Fatty Liver Disease: Marker and Mediator of Molecular Mechanisms. Mædica 2020, 15 (4), 513–516. 10.26574/maedica.2020.15.4.513.

(40) Martin-Armas, M.; Simon-Santamaria, J.; Pettersen, I.; Moens, U.; Smedsrød, B.; Sveinbjørnsson, B. Toll-like Receptor 9 (TLR9) Is Present in Murine Liver Sinusoidal Endothelial Cells (LSECs) and Mediates the Effect of CpG-Oligonucleotides. J. Hepatol. 2006, 44 (5), 939–946. 10.1016/j.jhep.2005.09.020.

(41) Akilesh, H. M.; Buechler, M. B.; Duggan, J. M.; Hahn, W. O.; Matta, B.; Sun, X.; Gessay, G.; Whalen, E.; Mason, M.; Presnell, S. R.; Elkon, K. B.; Lacy-Hulbert, A.; Barnes, B. J.; Pepper, M.; Hamerman, J. A. Chronic TLR7 and TLR9 Signaling Drives Anemia via Differentiation of Specialized Hemophagocytes. Science 2019, 363 (6423), eaao5213. 10.1126/science.aao5213.

(42) Liu, W.; Östberg, N.; Yalcinkaya, M.; Dou, H.; Endo-Umeda, K.; Tang, Y.; Hou, X.; Xiao, T.; Fidler, T. P.; Abramowicz, S.; Yang, Y.-G.; Soehnlein, O.; Tall, A. R.; Wang, N. Erythroid Lineage Jak2V617F Expression Promotes Atherosclerosis through Erythrophagocytosis and Macrophage Ferroptosis. J. Clin. Invest. 2022, 132 (13), e155724. 10.1172/JCI155724.

(43) Dupuis, L.; Chauvet, M.; Bourdelier, E.; Dussiot, M.; Belmatoug, N.; Le Van Kim, C.; Chêne, A.; Franco, M. Phagocytosis of Erythrocytes from Gaucher Patients Induces Phenotypic Modifications in Macrophages, Driving Them toward Gaucher Cells. Int. J. Mol. Sci. 2022, 23 (14), 7640. 10.3390/ijms23147640.

(44) Papadopoulos, C. Erythrocyte Glucotoxicity Results in Vascular Inflammation. Endocr. Metab. Immune Disord. Drug Targets 2022, 22 (9), 901–903. 10.2174/1871530322666220430013334.

(45) Papadopoulos, C.; Tentes, I.; Anagnostopoulos, K. Lipotoxicity Disrupts Erythrocyte Function: A Perspective. Cardiovasc. Hematol. Disord. Drug Targets. 2021, 21 (2), 91– 94. 10.2174/1871529X21666210719125728.

(46) Papadopoulos, C.; Tentes, I.; Anagnostopoulos, K. Molecular Interactions between Erythrocytes and the Endocrine System. Maedica 2021, 16 (3), 489–492. 10.26574/maedica.2020.16.3.489.

(47) Hotz, M. J.; Qing, D.; Shashaty, M. G. S.; Zhang, P.; Faust, H.; Sondheimer, N.; Rivella, S.; Worthen, G. S.; Mangalmurti, N. S. Red Blood Cells Homeostatically Bind Mitochondrial DNA through TLR9 to Maintain Quiescence and to Prevent Lung Injury. Am. J. Respir. Crit. Care Med. 2018, 197 (4), 470–480. 10.1164/rccm.201706-1161OC.

(48) Dinkla, S.; Wessels, K.; Verdurmen, W. P. R.; Tomelleri, C.; Cluitmans, J. C. A.; Fransen, J.; Fuchs, B.; Schiller, J.; Joosten, I.; Brock, R.; Bosman, G. J. C. G. M. Functional Consequences of Sphingomyelinase-Induced Changes in Erythrocyte Membrane Structure. Cell Death Dis. 2012, 3 (10), e410. 10.1038/cddis.2012.143.

(49) Hägerstrand, H.; Mrówczyńska, L.; Salzer, U.; Prohaska; Michelsen, K. A. Curvature-Dependent Lateral Distribution of Raft Markers in the Human Erythrocyte Membrane. Mol. Membr. Biol. 2006, 23 (3), 277–288.

(50) Terlizzi, M.; Colarusso, C.; Ferraro, G.; Monti, M. C.; Cerqua, I.; Roviezzo, F.; Pinto, A.; Sorrentino, R. Sphingosine-1-Phosphate Contributes to TLR9-Induced TNF-α Release in Lung Tumor Cells. Cell. Physiol. Biochem. Int. J. Exp. Cell. Physiol. Biochem. Pharmacol. 2021, 55 (2), 222–234. 10.33594/000000361.

(51) Devyatkin, A. A.; Revin, V. V.; Yudanov, M. A.; Kozlova, O. V.; Samuilov, V. D. Effect of Hydrogen Peroxide on Ejection of Cell Nucleus from Pigeon Erythrocytes and State of Membrane Lipids. Bull. Exp. Biol. Med. 2006, 141 (2), 261–264. 10.1007/s10517-006-0144-x.

(52) Arendt, B. M.; Ma, D. W. L.; Simons, B.; Noureldin, S. A.; Therapondos, G.; Guindi, M.; Sherman, M.; Allard, J. P. Nonalcoholic Fatty Liver Disease Is Associated with Lower Hepatic and Erythrocyte Ratios of Phosphatidylcholine to Phosphatidylethanolamine. Appl. Physiol. Nutr. Metab. Physiol. Appl. Nutr. Metab. 2013, 38 (3), 334–340. 10.1139/apnm-2012-0261.

(53) Dinkla, S.; van Eijk, L. T.; Fuchs, B.; Schiller, J.; Joosten, I.; Brock, R.; Pickkers, P.; Bosman, G. J. C. G. M. Inflammation-Associated Changes in Lipid Composition and the Organization of the Erythrocyte Membrane. BBA Clin. 2016, 5, 186–192. 10.1016/j.bbacli.2016.03.007.

(54) Zhu, X.; Song, J.; Mar, M.-H.; Edwards, L. J.; Zeisel, S. H. Phosphatidylethanolamine N-Methyltransferase (PEMT) Knockout Mice Have Hepatic Steatosis and Abnormal Hepatic Choline Metabolite Concentrations despite Ingesting a Recommended Dietary Intake of Choline. Biochem. J. 2003, 370 (Pt 3), 987–993. 10.1042/BJ20021523.

(55) Li, Z.; Agellon, L. B.; Allen, T. M.; Umeda, M.; Jewell, L.; Mason, A.; Vance, D. E. The Ratio of Phosphatidylcholine to Phosphatidylethanolamine Influences Membrane Integrity and Steatohepatitis. Cell Metab. 2006, 3 (5), 321–331. 10.1016/j.cmet.2006.03.007.

(56) Niebergall, L. J.; Jacobs, R. L.; Chaba, T.; Vance, D. E. Phosphatidylcholine Protects against Steatosis in Mice but Not Non-Alcoholic Steatohepatitis. Biochim. Biophys. Acta 2011, 1811 (12), 1177–1185. 10.1016/j.bbalip.2011.06.021.

(57) Wan, S.; van der Veen, J. N.; Bakala N’Goma, J.-C.; Nelson, R. C.; Vance, D. E.; Jacobs, R. L. Hepatic PEMT Activity Mediates Liver Health, Weight Gain, and Insulin Resistance. FASEB J. Off. Publ. Fed. Am. Soc. Exp. Biol. 2019, 33 (10), 10986–10995. 10.1096/fj.201900679R.

(58) Gao, X.; van der Veen, J. N.; Vance, J. E.; Thiesen, A.; Vance, D. E.; Jacobs, R. L. Lack of Phosphatidylethanolamine N-Methyltransferase Alters Hepatic Phospholipid Composition and Induces Endoplasmic Reticulum Stress. Biochim. Biophys. Acta 2015, 1852 (12), 2689–2699. 10.1016/j.bbadis.2015.09.006.

(59) Gonçalves, I. O.; Maciel, E.; Passos, E.; Torrella, J. R.; Rizo, D.; Viscor, G.; Rocha-Rodrigues, S.; Santos-Alves, E.; Domingues, M. R.; Oliveira, P. J.; Ascensão, A.; Magalhães, J. Exercise Alters Liver Mitochondria Phospholipidomic Profile and Mitochondrial Activity in Non-Alcoholic Steatohepatitis. Int. J. Biochem. Cell Biol. 2014, 54, 163–173. 10.1016/j.biocel.2014.07.011.

(60) Shama, S.; Jang, H.; Wang, X.; Zhang, Y.; Shahin, N. N.; Motawi, T. K.; Kim, S.; Gawrieh, S.; Liu, W. Phosphatidylethanolamines Are Associated with Nonalcoholic Fatty Liver Disease (NAFLD) in Obese Adults and Induce Liver Cell Metabolic Perturbations and Hepatic Stellate Cell Activation. Int. J. Mol. Sci. 2023, 24 (2), 1034. 10.3390/ijms24021034.

(61) Kanto, T.; Hayashi, N.; Takehara, T.; Katayama, K.; Kato, M.; Akiyama, M.; Kasahara, A.; Fusamoto, H.; Kamada, T. Low Expression of Erythrocyte Complement Receptor Type 1 in Chronic Hepatitis C Patients. J. Med. Virol. 1996, 50 (2), 126–134. 10.1002/(SICI)1096-9071(199610)50:2<126::AID-JMV5>3.0.CO;2-C.

(62) Burger, P.; Hilarius-Stokman, P.; de Korte, D.; van den Berg, T. K.; van Bruggen, R. CD47 Functions as a Molecular Switch for Erythrocyte Phagocytosis. Blood 2012, 119 (23), 5512–5521. 10.1182/blood-2011-10-386805.

(63) Gwag, T.; Ma, E.; Zhou, C.; Wang, S. Anti-CD47 Antibody Treatment Attenuates Liver Inflammation and Fibrosis in Experimental Non-Alcoholic Steatohepatitis Models. Liver Int. 2022, 42 (4), 829–841. 10.1111/liv.15182.

(64) Kirovski, G.; Dorn, C.; Huber, H.; Moleda, L.; Niessen, C.; Wobser, H.; Schacherer, D.; Buechler, C.; Wiest, R.; Hellerbrand, C. Elevated Systemic Monocyte Chemoattractrant Protein-1 in Hepatic Steatosis without Significant Hepatic Inflammation. Exp. Mol. Pathol. 2011, 91 (3), 780–783. 10.1016/j.yexmp.2011.08.001.

(65) Gäbele, E.; Mühlbauer, M.; Dorn, C.; Weiss, T. S.; Froh, M.; Schnabl, B.; Wiest, R.; Schölmerich, J.; Obermeier, F.; Hellerbrand, C. Role of TLR9 in Hepatic Stellate Cells and Experimental Liver Fibrosis. Biochem. Biophys. Res. Commun. 2008, 376 (2), 271–276. 10.1016/j.bbrc.2008.08.096.

(66) Lee, J.-G.; Lee, S.-H.; Park, D.-W.; Lee, S.-H.; Yoon, H.-S.; Chin, B.-R.; Kim, J.-H.; Kim, J.-R.; Baek, S.-H. Toll-like Receptor 9-Stimulated Monocyte Chemoattractant Protein-1 Is Mediated via JNK-Cytosolic Phospholipase A2-ROS Signaling. Cell. Signal. 2008, 20 (1), 105–111. 10.1016/j.cellsig.2007.09.003.

(67) Burger, P.; de Korte, D.; van den Berg, T. K.; van Bruggen, R. CD47 in Erythrocyte Ageing and Clearance – the Dutch Point of View. Transfus. Med. Hemotherapy 2012, 39 (5), 348–352. 10.1159/000342231.

(68) Gwag, T.; Reddy Mooli, R. G.; Li, D.; Lee, S.; Lee, E. Y.; Wang, S. Macrophage-Derived Thrombospondin 1 Promotes Obesity-Associated Non-Alcoholic Fatty Liver Disease. JHEP Rep. Innov. Hepatol. 2021, 3 (1), 100193. 10.1016/j.jhepr.2020.100193.

(69) Cui, W.; Maimaitiyiming, H.; Zhou, Q.; Norman, H.; Zhou, C.; Wang, S. Interaction of Thrombospondin1 and CD36 Contributes to Obesity-Associated Podocytopathy. Biochim. Biophys. Acta 2015, 1852 (7), 1323–1333. 10.1016/j.bbadis.2015.03.010.

(70) Tao, H.-C.; Chen, K.-X.; Wang, X.; Chen, B.; Zhao, W.-O.; Zheng, Y.; Yang, Y.-G. CD47 Deficiency in Mice Exacerbates Chronic Fatty Diet-Induced Steatohepatitis Through Its Role in Regulating Hepatic Inflammation and Lipid Metabolism. Front. Immunol. 2020, 11.

