## Supplementary figures and images for "Erythrocyte TLR9 is upregulated in metabolic associated fatty liver disease, and is linked to an inflammatory immunometabolic signature"

### Supplemental figure 1

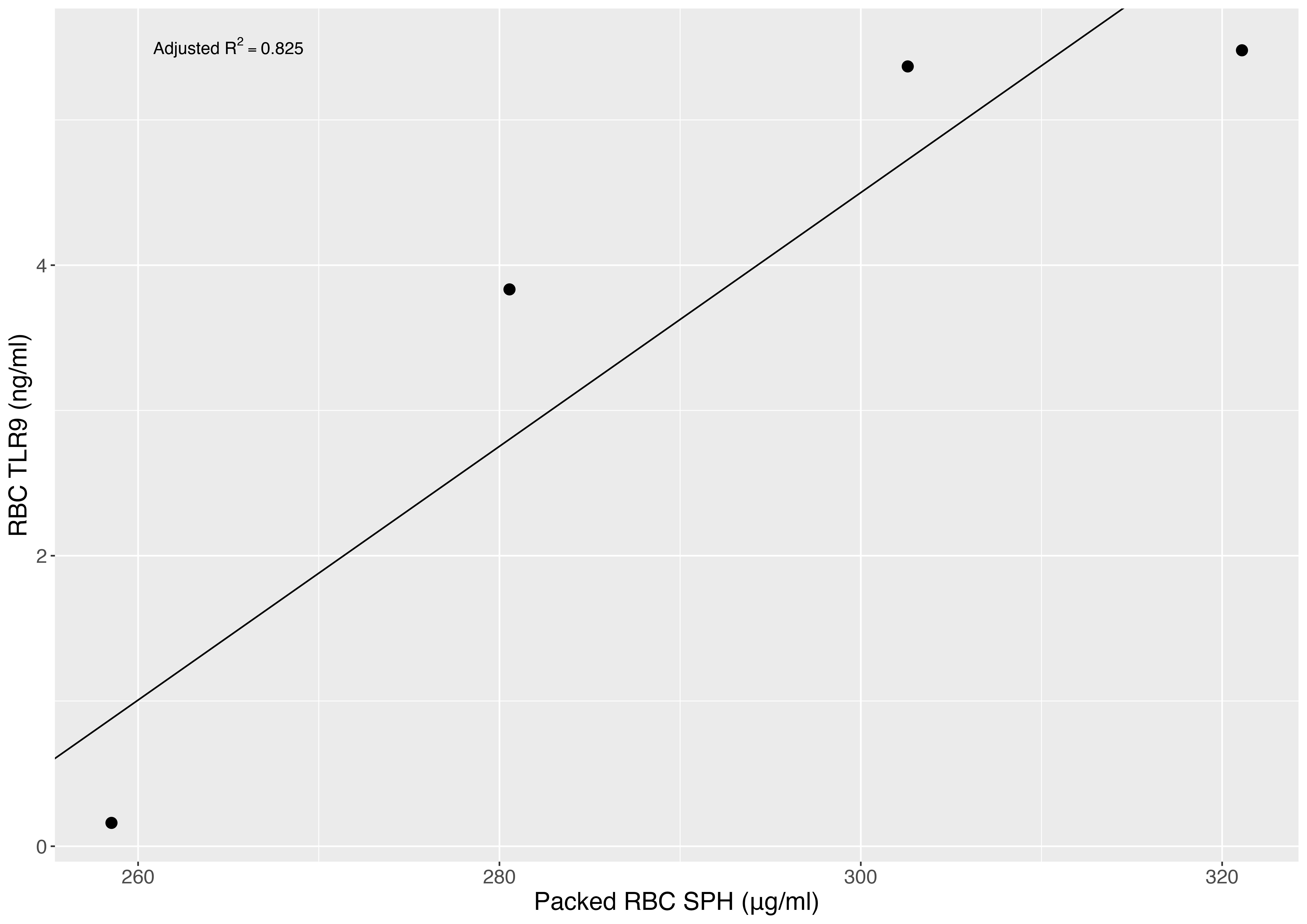
